# Haxe as a Swiss knife for bioinformatic applications: the SeqPHASE case story

**DOI:** 10.1101/2023.10.15.562406

**Authors:** Yann Spöri, Jean-François Flot

## Abstract

Haxe is a general purpose, object-oriented programming language supporting syntactic macros. The Haxe compiler is well known for its ability to translate the source code of Haxe programs into the source code of a variety of other programming languages including Java, C++, JavaScript and Python. Although Haxe is becoming more and more used for a variety of purposes, including games, it has not yet attracted much attention from bioinformaticians. This is surprising, as Haxe allows generating different versions of the same program (e.g. a graphical user interface version in JavaScript running in a web browser for beginners and a command-line version in C++ or Python for increased performance) while maintaining a single code, a feature that should be of interest for many bioinformatic applications. To demonstrate the usefulness of Haxe in bioinformatics, we present here the case story of the program SeqPHASE, written originally in Perl (with a CGI version running on a server) and published in 2010. As Perl+CGI is not desirable anymore for security purposes, we decided to rewrite the SeqPHASE program in Haxe and to host it at Github Pages (https://eeg-ebe.github.io/SeqPHASE), thereby alleviating the need to configure and maintain a dedicated server. Using SeqPHASE as an example, we discuss the advantages and disadvantages of Haxe’s source code conversion functionality when it comes to implementing bioinformatic software.

## 3 Introduction

Few biologists are proficient in using command-line tools [1]. As a result, bioinformatic software needs to be usable without the need to open a terminal. Nevertheless, some computer-savvy users prefer interacting with a tool via a command-line interface [2] (CLI) or need a command-line version to integrate the tool into a pipeline such as Galaxy [3–5]. Thus in practice most bioinformatic programs require two interfaces - a graphical user interface (GUI) and a CLI. Even though it is pretty straightforward to program a CLI, adding a GUI to a program can be trickier. GUIs can either be provided in form of a standalone application or via an external program such as a web browser (e.g. Chrome, Firefox, Edge or Safari). Although specialized toolkits such as Swing, SWT or FX for Java, or Flutter for Dart/C++ allow the creation of platform-independent standalone GUIs, this solution requires installing and maintaining the corresponding piece of software. By contrast, a web browser is preinstalled on most operation systems and is therefore more practical for biologists to use.

Historically, embedding code into a website was usually done using Java Applets or Flash applications with the Netscape Plugin Application Programming Interface [6]. However, newer web browsers do not allow this integration anymore due to security concerns [7]. Instead, modern browsers only allow the interpretation of a particular set of programming languages, namely WebAssembly [8, 9] and ECMAScript (with its better known dialect JavaScript) [10]. Due to this limitation, most programming languages cannot directly execute code inside a web browser. Thus, when programmers want to avoid maintaining two distinct versions of the same software (e.g. one written in JavaScript and the other one in Python), there are three possible ways to write a program that can be run both using a CLI and via a GUI running in a web browser:

- writing the whole program in JavaScript, then using e.g. NodeJS (https://nodejs.org) to execute the JavaScript program in a terminal environment. However, high-level programming constructs such as classes are rather awkward to use in JavaScript, except with the help of a scripting language such as CoffeeScript [11];
- writing a program with a CLI that runs on a web server. The GUI can then communicate with the program running on the web server and visualize its results (e.g. via BioJS [12]). Many scripting language such as PHP, Perl, Python or Ruby support communication via the common gateway interface (CGI)[13]. Nevertheless, setting up and maintaining such a dedicated public server can be time and resource-costly. It also requires users’ data to be sent over to the server via internet, which can be a problem in case of large and/or sensitive datasets;
- using a programming language that supports not only the interpretation of converting the source code into a binary program but also allows the conversion of the source code into the source code of various other programming languages (a process variously called “trans-compiling”, “transpiling” or “cross-compiling” depending on authors, and that is perhaps better described as “source-to-source translation”). A JavaScript version of the code can then run inside a web browser while another version of the program (e.g. in Python) can be used in a terminal environment.

One example of programming language enabling the latter approach is Haxe, a general purpose, highly versatile object-oriented programming language [14]. Although other source-to-source compilers exist, such as Dafny [15], Haxe is the most widely used among them. The source code of Haxe programs can be converted into the source code of a variety of other languages including Java, C++, JavaScript and Python [16]. Using Haxe, it is thus relatively easy to create multiple versions of the same tool - e.g. a JavaScript version that runs inside a web browser and a Python version that can run as a terminal application. Furthermore, Haxe supports sophisticated programming paradigms such as syntactic macros [17], making it an excellent choice for writing bioinformatic applications.

To illustrate the usefulness of Haxe as a Swiss knife for bioinformatic applications, we tell here the story of how we used Haxe to revive an old but very useful piece of code, SeqPHASE [18], originally written in Perl + CGI code, by reimplementing it in Haxe then converting the Haxe code into JavaScript (for the graphical version) and Python (for the command-line one).

## 4 Material and methods

SeqPHASE was written in 2010 to address a pressing practical issue in population genetics: how to input FASTA files into PHASE [19] (a program originally written to infer the most probable pairs of haplotypes in a population of diploid organisms for which genotypes have been determined, but designed for microsatellites and not DNA sequences), and how to the output of PHASE back into FASTA [18].

Until that point, a tool allowing this conversion existed as part of the Windows program DNAsp [20, 21], but with several severe shortcomings [18], and conversion was impossible on other operating systems such as Linux or MacOS. Because it filled an important need in the biological community, SeqPHASE was immediately adopted and cited a relatively large number of times (more than 500 times since published in 2010).

However, its implementation in Perl + CGI posed important security issues, and at some point university computer infrastructure administrators became very reluctant to host this piece of code on public servers as it could offer an entrance point to hackers. Out of this necessity, it was therefore decided to reimplement SeqPHASE completely; this time in Haxe as we needed to provide both a user-friendly web browser tool and a command-line version for use in pipelines and on computer clusters.

For the sake of simplicity and cost-saving, a choice was made to host the Haxe source code on GitHub (https://github.com/eeg-ebe/SeqPHASE). Compiling the Haxe part of the code into JavaScript resulted in a fully functional web browser tool accessible via GitHub Pages (https://eeg-ebe.github.io/SeqPHASE), whereas compiling the same Haxe code into Python produced a cross-platform Python script (with a CLI) made available for download on the same website (https://eeg-ebe.github.io/SeqPHASE/download.html).

## 5 Results

The source code of the reimplemented SeqPHASE program is available at https://github.com/eeg-ebe/SeqPHASE and licensed under the Apache 2.0 license. A web page allowing the user to run and/or download the program is available on GitHub Pages https://eeg-ebe.github.io/SeqPHASE.

The reimplementation consists of the following files:

- a series of static HTML pages including a menu page, a FAQ page and so on;
- two dynamic web pages where users can run respectively for the first (FASTA to PHASE) and second step (PHASE to FASTA) of the SeqPHASE program. The web pages directly calls the corresponding JavaScript code;
- a download page that allows users to obtain a zipped archive of the GUI version of the program (once unzipped, users can run SeqPHASE offline inside a web browser by double-clicking the index.html file found inside that archive) as well as Python command-line versions of the two steps of SeqPHASE.

During the reimplementation process we discovered small bugs in the original code and corrected them, namely:

- for Step 1 (FASTA to PHASE), every sequence in the inputted FASTA files should have the same length, but it was possible to bypass this check by ordering the sequences by decreasing length;
- for Step 2 (PHASE to FASTA), the sequences of individuals were sorted apart if the name of one sequence was a full prefix of a longer name of another sequence (e.g. the name of sample1a and sample1b would be sorted apart if there was another sequence with the name sample1abc1a).

Although it is unlikely that these bugs had an impact on the accuracy of the results, it sometimes resulted in the program running on inputs that contained errors, instead of reporting those errors.

## 6 Time usage

In order to analyze the runtime usage of the SeqPHASE programs, we used Haxe to create different program versions with C++, Lua, Neko, Python, JavaScript (NodeJS) and Java as target languages (Table 1): one program version where Haxe’s dead code elimination algorithm was turned on, one version where Haxe’s dead code elimination algorithm was turned off and one version where Haxe’s dead code elimination algorithm was limited to classes in the Haxe standard library. This process resulted in 18 executables (= six target programming languages ^*^ three dead code elimination strategies) for the FASTA to PHASE conversion process as well as 18 other executables for the PHASE to FASTA conversion process.

**Table 1:** Versions of the different compilers/interpreters used for our benchmark.

|  | Linux | MacOS |
| --- | --- | --- |
| haxe | 4.2.4 | 4.2.4 |
| lua | 5.1.5 | 5.4.6 |
| neko | 2.3.0 | 2.3.0 |
| python | 3.10.12 | 3.9.10 |
| perl | 5.34.0 | 5.30.3 |
| java | 11.0.22 | 15.0.1 |
| nodejs | 12.22.9 | 20.11.0 |
| c++ | g++ 11.4.0 | clang 1300.0.27.3 |

Twenty FASTA data files of variable sizes were generated using SimCoal [22]. We then launched the 18 programs as well as the two original Perl programs 120 times on each of these 20 datasets and measured the corresponding time usages (compare figure 6 and table 3). To evaluate startup times, we also created Hello World executables by source-to-source translating a Haxe Hello World program to the different target programming languages, as well as writing a Perl Hello World program (Figure 6 and Table 2).

**Table 2:** Time usage (in ms) of the Hello World program on Linux and Mac. The table lists the minimum, maximum, standard deviation, average and median time of the Perl, C++, Lua, Python, Neko, NodeJS and Java Hello World programs.

| Program | Linux |  |  |  |  | Mac |  |  |  |  |
| --- | --- | --- | --- | --- | --- | --- | --- | --- | --- | --- |
| | Min | Max | $\sigma$ | Avg | Med | Min | Max | $\sigma$ | Avg | Med |
| lua | 6.1 | 10.6 | 0.9 | 7.6 | 7.4 | 7.2 | 11.0 | 1.3 | 9.0 | 9.6 |
| neko | 8.5 | 26.0 | 3.5 | 13.9 | 13.0 | 9.8 | 13.1 | 1.0 | 11.0 | 10.7 |
| python | 32.1 | 49.0 | 3.2 | 37.5 | 37.2 | 40.7 | 61.8 | 6.6 | 51.6 | 53.6 |
| perl | 5.2 | 10.5 | 0.9 | 6.3 | 6.14 | 7.7 | 12.1 | 1.3 | 9.2 | 9.0 |
| java | 80.0 | 136.0 | 8.6 | 94.0 | 93.6 | 68.7 | 85.2 | 4.8 | 73.9 | 73.9 |
| nodejs | 75.9 | 123.2 | 7.1 | 87.0 | 86.5 | 43.7 | 53.9 | 3.7 | 47.7 | 47.2 |
| c++ | 5.6 | 15.9 | 1.4 | 6.9 | 6.6 | 7.2 | 15.0 | 2.2 | 9.3 | 9.0 |

**Table 3:**
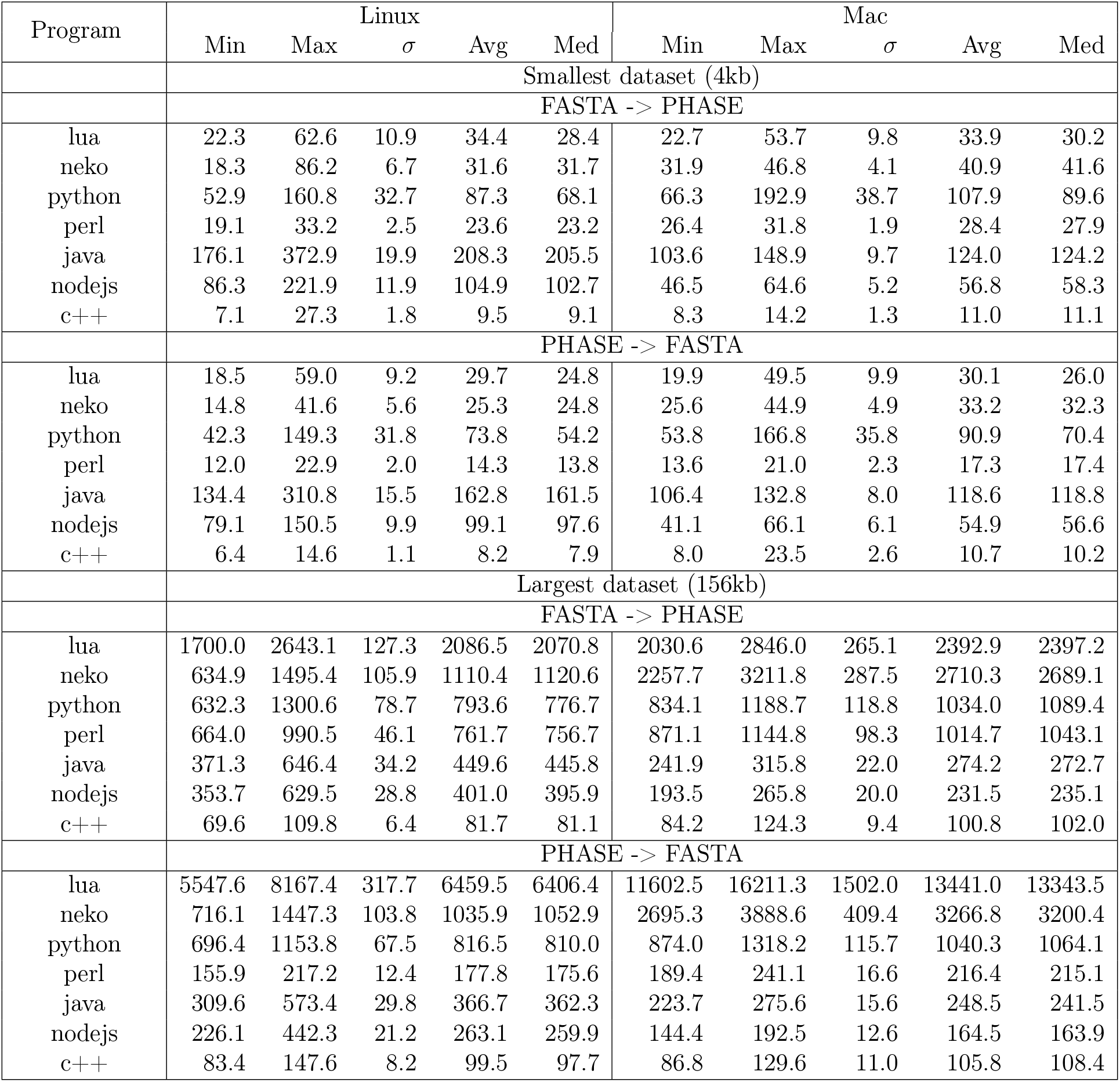
Time usage (in ms) for different dataset sizes for the two different steps of the SeqPHASE on Linux and Mac. The table lists the minimum, maximum, standard deviation, average and median time of the Perl, C++, Lua, Python, Neko, NodeJS and Java SeqPHASE programs.

All datasets and source codes used in this benchmark are available at https://github.com/eeg-ebe/SeqPHASE_time_mesurements.

**Figure 1:**
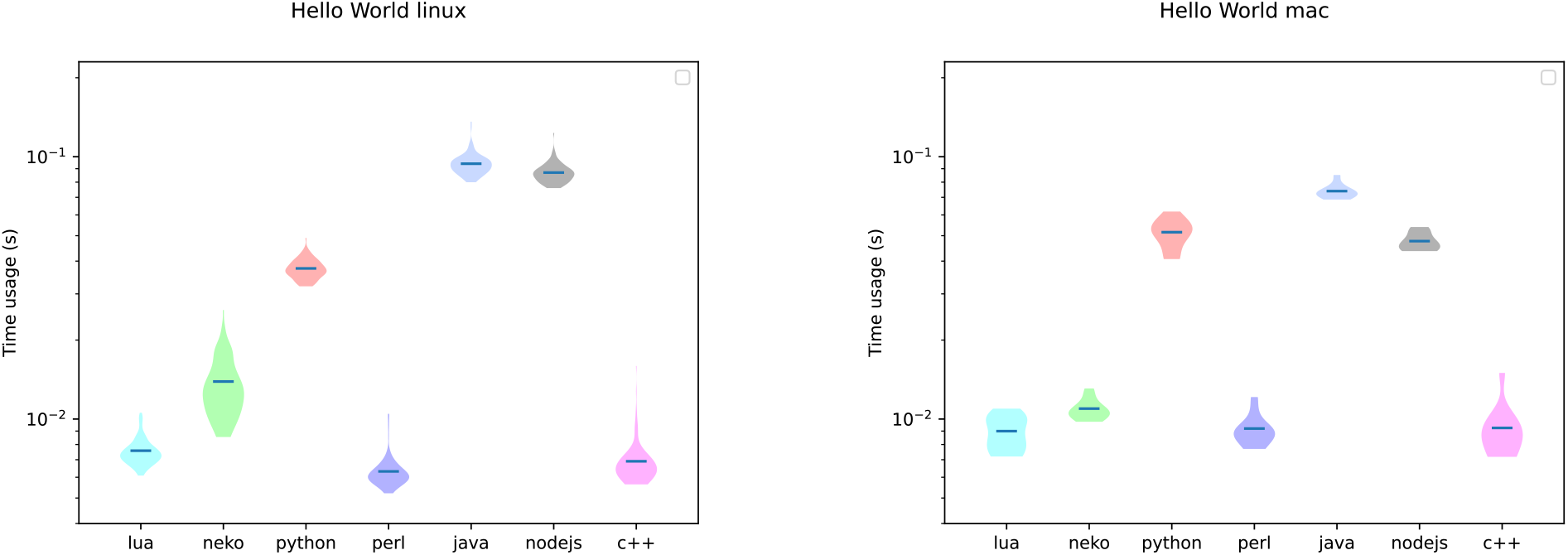
For this plot, each of the 7 Hello World executables (1 Perl executable + 6 executables created by source-to-source translation of the Haxe Hello World source code into the corresponding target language) were launched 120 times. The left panel shows the time usage of the Hello World programs on a Linux machine while the right panel shows the corresponding analysis on a computer running the MacOS operating system. The minimum, maximum, standard deviation, average and median of these runs for the different program versions and conversions are listed in Table 2.

When comparing the average calculation time needed to convert a FASTA file to a PHASE input file or a PHASE output file back to a FASTA file, the C++ version greatly outperformed all other versions of the SeqPHASE program, followed by - for small datasets - the Python / Neko / Lua versions. However, for larger datasets the overhead of starting a virtual machine with a long startup time can pay off. In that case, the NodeJS and / or Java version may outperform every other versions except for the C++ version.

**Figure 2:**
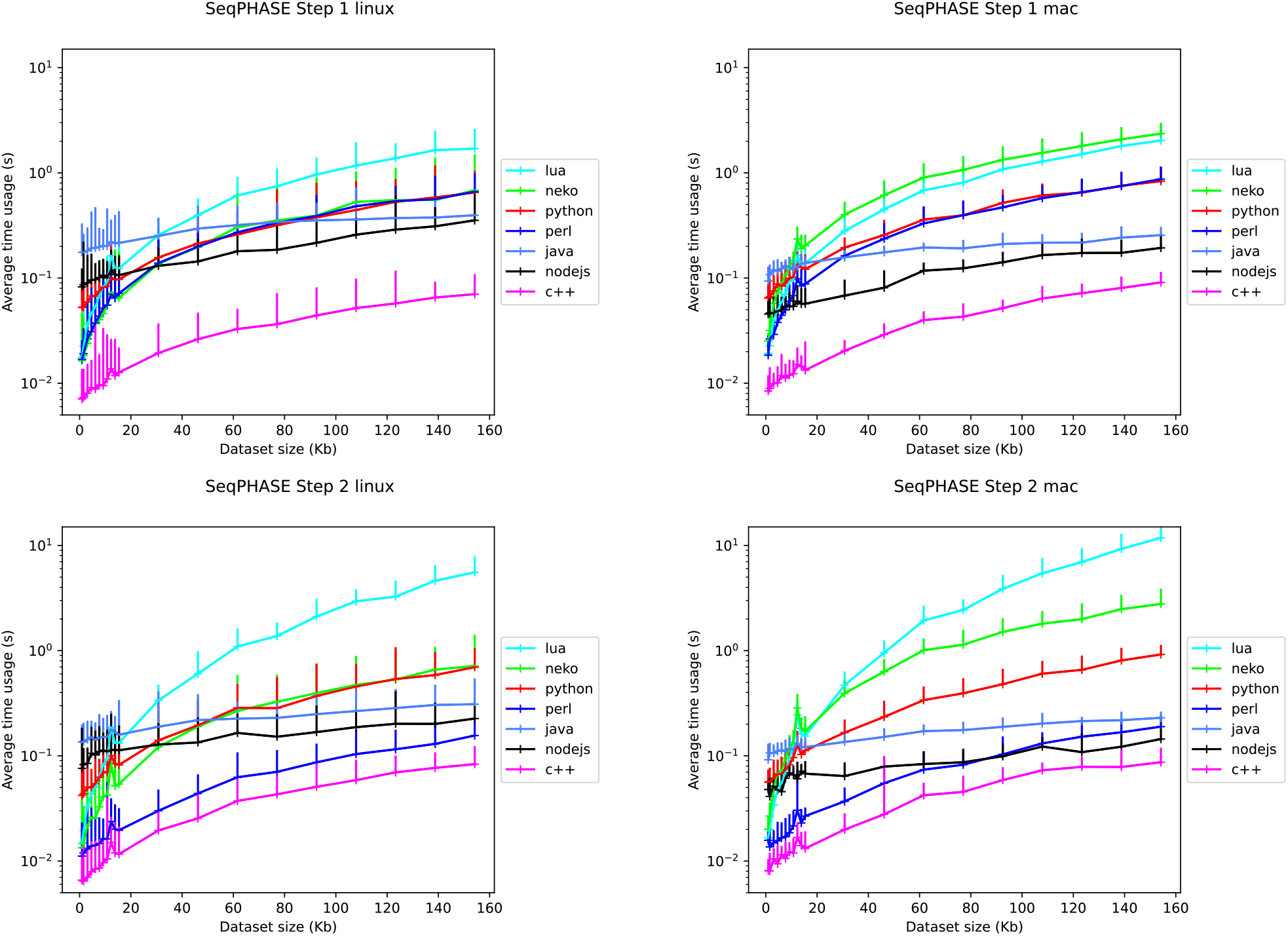
Measurement of the average time usage the SeqPHASE program takes to convert different dataset sizes. The top panels show the calculation time needed for the conversion of the FASTA file into the PHASE input file format, while the bottom panels show the conversion of the PHASE output file format to FASTA file format, for Linux (left) and for MacOS (right). The error bars indicate the measured minimum and the maximum calculation time measured. The minimum, maximum, standard deviation, average and median of these runs for the different program versions and conversions are listed in Table 3.

Although the C++ version was the fastest we decided to not provide it for download as we found it challenging to compile a C++ program that runs on all computers. Instead we opted for the Python version because Python is a well-known programming language already installed on most computers and because speed is not that much of an issue when it comes to small, straightforward programs that are running in less then 100ms, such as SeqPHASE. However, in case high-performance is needed, building a C++ version for a particular target computer with platform-specific code optimization would be the best option.

Even though this assumption still needs to be verified, we believe that Chrome and NodeJS would need, on average, the same amount of time to execute the SeqPHASE program since both programs are using the same JavaScript engine (namely, V8 [23]).

When comparing the dead code elimination strategies for the different target programming languages, we did not observe any differences in the execution times of the versions created when the dead code elimination strategy was applied to the full source code or was limited to the Haxe standard library (the default option). However the versions created without any dead code elimination were significantly slower for most programming languages, except for C++, Neko and Java for which no difference was observed.

## 7 Discussion

Through the example of SeqPHASE’s reimplementation, we illustrate how Haxe coding is a valuable, yet still underused approach for bioinformaticians to make programs available online to a large audience including both biologists with no command-line proficiency and computer scientists requiring command-line tools to run on computing clusters. The SeqPHASE website can be used as a template for users wishing to experiment with using Haxe as a valuable alternative to CGI apps, for instance.

Although we chose a fairly simple program to illustrate and benchmark our proposed approach, another more complex example of a bioinformatic tool we reimplemented in Haxe in replacement for a previous perl+CGI version is https://eeg-ebe.github.io/Champuru [24, 25]. Moreover, three further tools we directly implemented in Haxe are available online at https://eeg-ebe.github.io/HaplowebMaker and https://eeg-ebe.github.io/CoMa [26] as well as https://eeg-ebe.github.io/KoT [27].

Since websites are running inside a sandboxed web browser, tools that run inside websites are also advantageous for users that do not want to install a particular software locally due to security concerns. However, compared to Perl, Python and other programming languages traditionally used in bioinformatics, Haxe suffers from certain drawbacks: (1) the current unavailability of a “BioHaxe” library of functions facilitating the import and processing of biological data. It is our hope to provide such a library in the future; (2) debugging a particular Haxe program may require to take a look and understand its translation into the target language, which necessitates some understanding of this language (in addition to knowing Haxe); (3) although Haxe supports the languages most often used in bioinformatics (Python, C++, JavaScript, Java), some other languages such as Perl and R are not supported yet.

## 8 Author Biographies

**Yann Spöri** is a PhD student at the Department of Organismal Biology of the Université libre de Bruxelles (ULB). His doctoral research deals with improving computational methods for species delimitation. **Jean-François Flot** is a professor at the Department of Organismal Biology of ULB and a member of the Interuniversity Insitute of Bioinfomatics in Brussels – (IB)^2^. His research group ‘Ecological and Evolutionary Genomics’ investigates how the ecology of organisms influences the evolution of their genomes.

## 9 Key Points

- The programming language Haxe allows designers of bioinformatic applications to maintain a single code for both command-line and graphical-user-interface versions of their program.
- This Haxe source code can then be compiled into various languages such as C++, JavaScript or Python.
- As a case study to illustrate Haxe’s usefulness for bioinformatics, we reimplemented in Haxe the previously published program SeqPHASE in Haxe (originally written in Perl) and compared the performances of translated C++, Lua, Python, Neko, NodeJS and Java versions.

